# Egg colour mimicry in the common cuckoo results from coevolutionary alternation, not coevolutionary arms races

**DOI:** 10.64898/2025.12.01.691685

**Authors:** Daniel Hanley, David Kepplinger, Ariba Nasir, Peter Samaš, Juliana Villa, João Vitor de Alcantara Viana

## Abstract

The exquisite eggshell mimicry found in many cuckoo-host systems has been the textbook example of a coevolutionary arms race^1^. In this system, cuckoos lay their eggs in their hosts’ nests leaving these foster parents to rear their young^2^. In defence, most of these hosts have evolved the ability to recognize cuckoo eggs. It is long-assumed that these interactions select for ever-improving eggshell mimicry in cuckoo host-races laying distinct eggs that specifically mimic their host’s eggshell, regardless of its particular colour^3,4^. We challenge this paradigm and demonstrate that coevolutionary arms races and egg colour mimicry are the exception to the rule. Here, using eggshell reflectance spectra from 39 host species and the specific cuckoo eggs found in their nests, we show that eggshell colour mimicry is very uncommon even among frequent hosts. Although cuckoo eggs display considerable variation in eggshell colour, their hosts’ eggshell colours occupy 6 times their colour volume. As a result, mimicry is only achieved for a small subset of the egg colour gamut. Furthermore, cuckoo host-races rarely lay eggs that match their hosts’ eggs better than those of other potential hosts or that are distinct from those laid by other cuckoo host-races. These patterns are inconsistent with the long-held assumption of coevolutionary arms races, but consistent with coevolutionary alternation that is defined by a process of continual host-switching. These findings open new avenues for research in a system that has long served as a model for studying coevolutionary processes.

## Main

Coevolved eggshell mimicry of obligate avian brood parasites^1,2^, such as the common cuckoo *Cuculus canorus* (hereafter, cuckoo), has long fascinated scientists^5^ and has been especially vital in our understanding of coevolutionary processes^3,6,7^. In particular, the apparent mimicry of cuckoo eggs to a wide array of hosts has provided a clear example of coevolutionary arms races^6,8^. Under this presumed process of reciprocal selection pressure, hosts evolve the ability to recognize and reject parasitic eggs, which in turn selects for improved egg mimicry in their cuckoos^6^. This instigates a cycle of ever-improving recognition on part of the host and egg mimicry on part of the cuckoo. Since most of the cuckoo’s hosts use eggshell colours and patterns to recognize cuckoo eggs, cuckoo eggshell mimicry tends to be better for these traits than for traits that hosts rarely use for recognition, such as size or shape^9^. By removing eggs that hosts perceive as different from their own, we have long assumed that hosts select for improved cuckoo eggshell mimicry^3,4,6,10^. This incredible example of a coevolutionary arms race can occur because although the cuckoo parasitizes a wide range of hosts^5,11^, individual cuckoo females tend to parasitize their natal hosts^12–15^ and they mostly inherit egg traits from their mothers^12–14,16^. Thus, host egg recognition selects for distinct cuckoo eggshell mimicry in the specific matrilineage of cuckoos that parasitizes them (hereafter host-race)^3,4,8^, while ever-improving eggshell mimicry reciprocally selects for hosts with greater egg recognition abilities^17,18^.

Most research on cuckoo eggshell colour mimicry has focused on host egg recognition (adaptation) and cuckoo eggshell morphology (counter-adaptation), as a framework for testing coevolutionary arms races^4,19^. However, coevolutionary arms races would result in numerous testable predictions. For example, if cuckoo eggshell colour mimicry coevolves via arms races, we predict that, given sufficient coevolutionary history and selection pressure^7^, cuckoos (i) could evolve egg colour mimicry to any host, regardless of their egg colour^20–24^. As such, they should lay eggs that (ii) specifically match those laid by their particular host species^3^. Finally, cuckoo host-races that specialize on particular host species^15^ should (iii) converge on similar eggshell appearances to those targeting the same host species. Surprisingly, we still lack conclusive support for these basic predictions. Therefore, we now test these using eggshell reflectance spectra from host and cuckoo eggs from 468 parasitized host clutches spanning 39 hosts species, incorporating new data and data from a classic study^4^ as a form of metareplication.

### Testing current dogma

Contrary to our expectations under coevolutionary arms races, we found that eggshell mimicry is directly related to host eggshell colour, such that some host eggshell colours are easier to match than others (Conditional R^2^=0.85, CI=[0.77, 0.90], *p* < 0.01; Fig. 1). The cuckoo’s eggshell colour mimicry is significantly worse for those parasitizing hosts with colourful eggs (either blueish or brownish; Fig. 1). By contrast eggshell mimicry is most refined for hosts that lay eggs ranging from white to light brown eggs (Fig. 1). In fact, the colour of individual cuckoo eggs that achieved the best eggshell colour mimicry were not significantly different than the colour of a pure white egg (Conditional R^2^=0.49, *p*=0.73; Fig. S1). This may be because all white eggshell colours are generated by calcium carbonate^25,26^, thus white (or very pale) eggs will inherently have visually similar colours; however, this explained relatively little of the variation in mimicry. A more plausible argument for the clear relationship between host eggshell colouration and cuckoo eggshell mimicry could be that cuckoos have just a few typical egg-types that do not specifically target all host species, suggesting a generalized parasitic strategy^27,28^.

**Figure 1.**
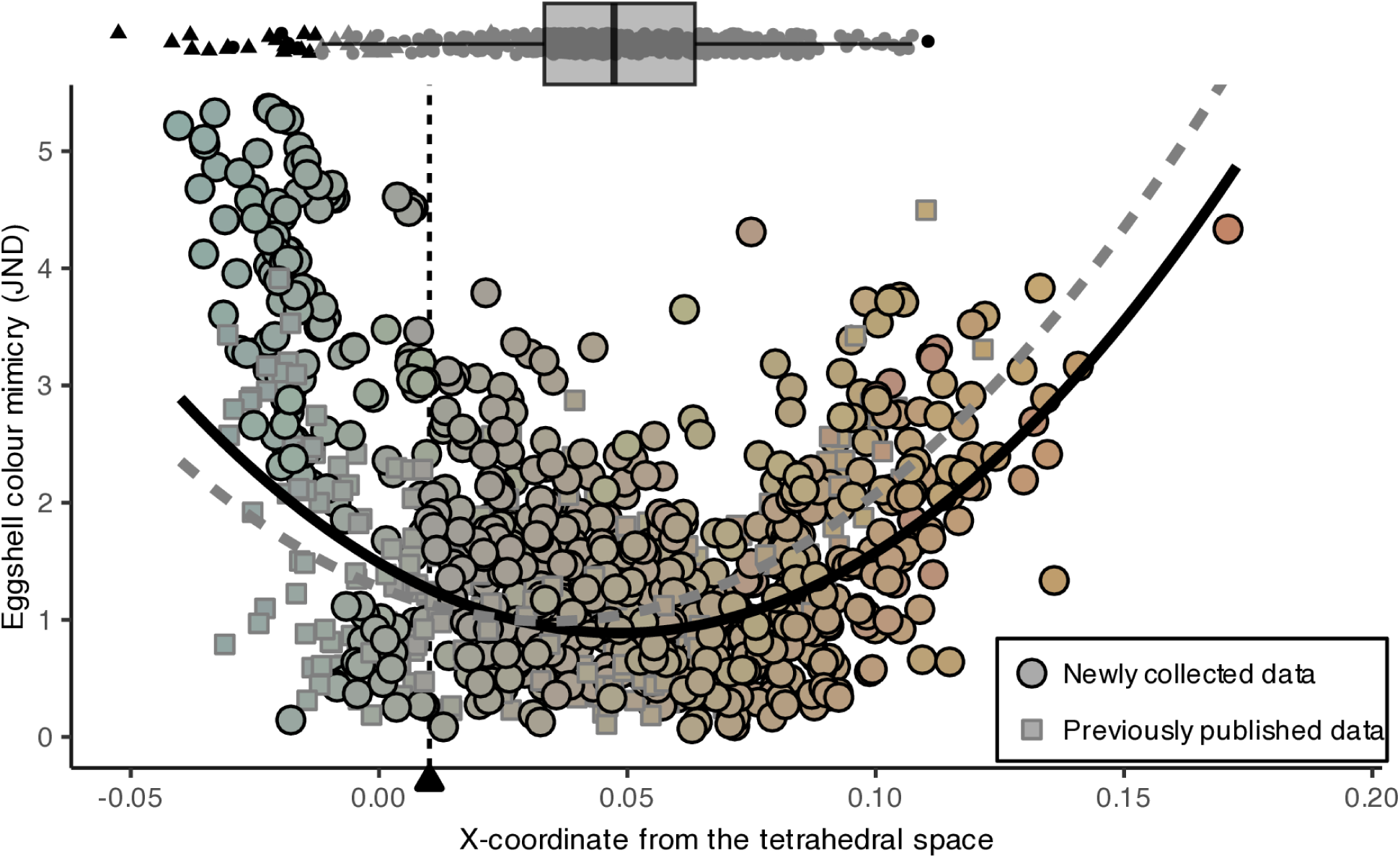
The eggshell colour mimicry (JND) achieved by 474 individual cuckoos to their specific hosts. Eggshells with lower values of eggshell mimicry represent better matches from the perspective of the cuckoo’s hosts. Here we plot both newly collected data (black circles) and data from previously published results^4^ (grey squares) from 39 host species. In this case the fit for the combined dataset (solid black line) is very similar to that from the previously published results (dashed blue line). The match in eggshell colour improves near the colours we expect for white and white speckled eggs (the ‘colour’ of a white mourning dove egg is indicated by a black triangle). We show a boxplot of all cuckoo eggshell colours above the scatterplot to illustrate the distribution of cuckoo eggshell colours relative to their hosts’ eggshell colours.

We found that cuckoo eggs were far less colourful than host eggs (Fig. 1). Specifically, across the cuckoo’s entire host community, hosts’ eggs covered more than 6 times the colour volume of the cuckoo’s eggs (*relative volume*=6.4, 95% CI=[3.5, 11.2], *p*<0.0001). Thus, hosts that lay more colourful eggs are not well matched because cuckoo eggshell colours were much less colourful and generally expressed colours from a relatively restricted range of the egg colour gamut (Fig. 1 & S2). That said, when compared to individual host species, cuckoo eggs display a much broader range of colours than the eggs of all of their host species (all *p*<0.0001; Fig. S3), except for the tree pipit *Anthus trivialis* that lays eggs with a comparable egg colour volume as the cuckoo’s eggs (*relative volume*=1.2, 95% CI=[0.4; 2.2], *p*=0.46; Fig. S3). These findings strongly suggest that cuckoo egg phenotypes have evolved in response to egg rejection or have evolved to be highly variable^29^, as seen in other systems^30^; however, they also demonstrate that cuckoo eggshell colours do not span the same range as the eggshell colours of their host community.

Under coevolutionary arms races, we also expect that cuckoo host-races should lay eggs that specifically match their hosts; however, we found that only three cuckoo host-races lay eggs with colours better matched to their specific host than other host species (Fig. 2; Table S1): the European robin *Erithacus rubecula* (R^2^=0.69, *p*=0.02), the yellow wagtail *Motacilla flava* (R^2^=0.70, *p*=0.04) and the Western Orphean warbler *Sylvia hortensis* (R^2^=0.95, *p*=0.03). The latter two host-races were rare and significance should be viewed with caution. In fact, of the 39 host races, only 29 host-races were common enough to calculate robust estimates of within host-race variability. Of these, only the host-race parasitizing the European robin specifically matched its host. All other cuckoo host-races were just as likely to produce a colour that matched their host as any other host species (Fig. 2; Table S1), which suggests evolutionary generalization instead of specific host-race specialization^27,31–33^. We found surprisingly little evidence of host specificity (i.e., cuckoo eggshell colours specifically targeting certain hosts), which implies that many host eggs may simply be similar in colour to the cuckoo’s egg, without any coevolved mimicry^34^; pure coincidence or convergence could easily be confused for mimicry^35^. In fact, the more limited egg colour gamut of cuckoo eggs suggests that this is plausible in many cases^28,36,37^, particularly for relatively unpigmented white eggshells^26^. By comparison, while host species rarely laid distinctly coloured eggs, the variation in eggshell colouration was always greater between than within host species (Fig. S4; Table S2). When comparing every parasitism event, cuckoos rarely targeted hosts that laid similar egg colours (Fig. S5). Overall, these findings suggest that cuckoos do not specifically mimic the colours of their current hosts’ eggs.

**Figure 2.**
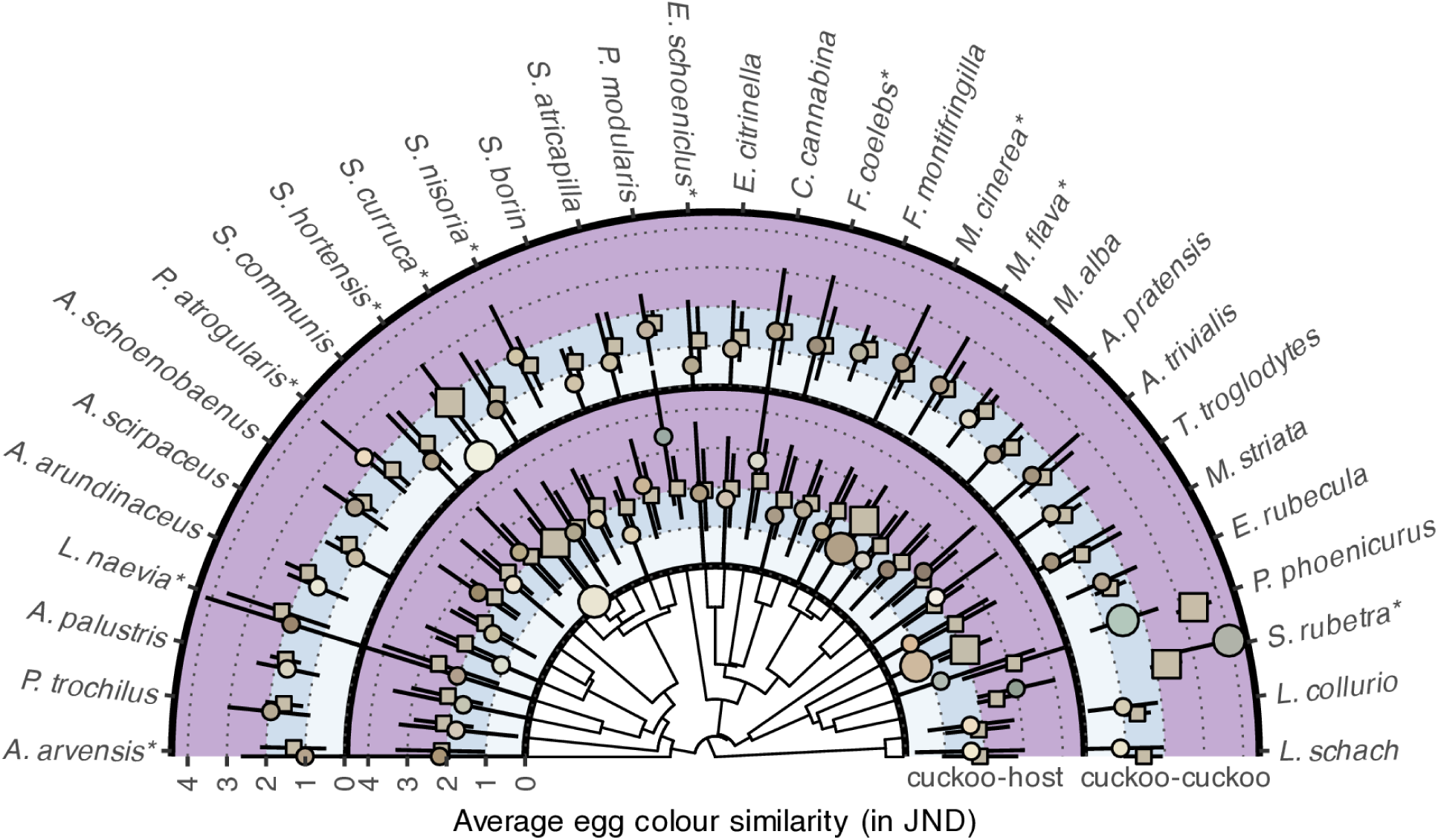
Average perceived difference in eggshell colour (in JND) among 32 cuckoo host-races and their respective hosts. These differences are depicted along the edges of a maximum clade credibility tree defining the relationships among the host species. The egg colour differences between cuckoo and host eggs are shown along the inner ring (labelled ‘cuckoo-host’), while the differences among cuckoo host-races are shown on the outer ring (labelled ‘cuckoo-cuckoo’). Egg colour differences between each cuckoo host-race and their specific host or among other cuckoos of the same host race are shown in circles on the inner and outer ring respectively. Egg colour differences between each cuckoo host-race and other hosts or other cuckoo host-races are shown in squares. All data are shown in mean with 95% confidence intervals. Bands of colour represent differences < 1 JND (white), ≤ 2 JND (pale blue), and > 2 JND (light purple). We have enlarged the points representing significant differences. We excluded 7 rare host-races for which we could not estimate within host variation because there were < 2 specimens, and use asterisks to indicate host-races with sparse sampling (≤ 3 specimens) that should be interpreted with care (Table S1 & S2). The circles are coloured based on the average colour for each host-race, while the squares are coloured as the average colour across all host-races.

Finally, we often assume that cuckoos form distinct host-races that lay eggs with colours visually distinct from those of other cuckoo host-races^15^; however, contrary to these expectations, we found that only two cuckoo host-races laid eggs with colours that were visually distinct from those laid by other cuckoo host-races (Fig. 2; Table S3). Specifically, these were the host-races parasitizing the common redstart *Phoenicurus phoenicurus* (R^2^=0.84, *p*<0.001) and the western Orphean warbler (R^2^=0.96, *p*=0.01). Interestingly, the cuckoos parasitizing the whinchat *Saxicola rubetra* laid eggs less similar in colour to cuckoos from its own host-race, yet more similar to those from other host-races (R^2^=0.07, *p*=0.01). However, both the whinchat and western Orphean warbler were rare host-races and these findings should be viewed with caution. By contrast, the cuckoo that parasitizes the common redstart *Phoenicurus phoenicurus* was the only common host-race that produced an egg that was distinct from those laid by other cuckoo host races. That said, the distinct blue-green eggshell phenotype of this host-race did not match its host significantly better than it matched the colour of eggs laid by other hosts (R^2^=0.94, *p*=0.08; Fig. 2; Table S1); the immaculate blue-green colour of the egg of this cuckoo host-race would provide a reasonable match to the blue-green of other hosts. In this case, the redstart-type cuckoo has a unique genetic mechanism that gives rise to its distinctive pale blue egg^38^. The redstart’s cavity nesting habits, coupled with its unique immaculate blue egg, may present an evolutionary trap for this host-race, which could explain why it is also genetically distinct^38^ and why its egg is unlike the other host-races (Fig. 2). Overall, eggshell colours among cuckoo host-races were largely not visually distinct, and any colour similarity between the eggs of cuckoo host-races could just as easily result from fortuitous host-switches as coevolved mimicry.

Coevolutionary arms races imply that cuckoos should evolve into distinct, host-specific, races that lay eggs mimicking the appearance of their hosts’ eggs, regardless of the hosts’ egg colours. Our results did not support these long-held assumptions.

### Complex host-parasite dynamics

Our findings suggest that the coevolutionary dynamics often attributed to cuckoos and their predominant hosts^18^ is overly simplistic. The strict dyads between current hosts and cuckoo host-races are probably far more fluid and cuckoo lineages likely frequently host switch^12,27,39^. Coevolutionary dynamics require detailed knowledge of their long-term interactions and of reciprocal evolutionary changes, which we largely lack for this system^7^. However, we do know that host usage can vary markedly over time^40,41^. In Japan, the cuckoo exploited a new host over the course of a half century^40^, yet has not evolved a mimetic egg^36^. Likewise, some cuckoo host-races have experienced dramatic declines in the UK (e.g., dunnock and robin host-races), while there have been simultaneous increases of other cuckoo host-races (e.g., the reed warbler-type) over a 50-year period^41^. Such boom and bust cycles, may be common for cuckoo host-races. The red-backed shrike *Lanius collurio* is the perfect example of a historically frequent host^42,43^ that experienced a dramatic decline in usage over the 20^th^ century, such that it is now a very rare host^44–46^. Despite this decline in parasitism rates, the red-backed shrike is still highly aggressive to cuckoos and has expert egg recognition abilities, which may suggest that this host ‘won’ the arms race forcing its cuckoo host-race on to other host species^47^ or alternatively that it drove its cuckoo host-race to extinction. Intriguingly, between 2022-2024 four cuckoo eggs (very distinct from those of the shrike) were found in red-backed shrike nests^48^, suggesting that a cuckoo host-race may be switching back to this once common host.

Cuckoo eggshell colours are mostly maternally inherited^16^, via mitochondrial DNA and W chromosome DNA^12,13,15,16,49,50^; however, these studies also uncover complex relationships among and between cuckoo host-races^12,51^. For example, mitochondrial haplotype networks for the reed warbler-, meadow pipit-, and dunnock-type cuckoo host-races likely result from frequent host switching^12^. By contrast, the redstart-type cuckoo diverged and has remained distinct from other European host-races^38^, which could explain why this host-race was the only host-race with a distinct eggshell colour (Fig. 2). This frequent alternation of hosts could easily explain why we found a general lack of specificity and distinctiveness of cuckoo eggshell colours. Interestingly, both genetic studies and a number of field studies suggest that cuckoos will either occasionally or regularly^27,52,53^ use alternative hosts. Thus individual female cuckoos are known to parasitize multiple host species^27,54^, such that each season siblings will be reared by different host species. Moreover, cuckoo eggshell colours may be evolutionary constrained; cuckoo host-races with eggshell phenotypes targeting extreme host phenotypes may be at a disadvantage because the matrilineal inheritance of background eggshell colour limits continued coevolution due to lack of potential for recombination^16,55^. This would reduce their opportunity to host-switch when their hosts adapt strong recognition abilities, which would leave cuckoo host-races that have evolved extreme eggshell colours at a greater risk of extinction. Additionally, recent research has found that male cuckoos contribute to the eggshell phenotypes through autosomal inheritance of eggshell patterns^16^, further limiting the potential to achieve perfect mimicry. In fact, specific mimicry could be challenging to evolve unless eggshell colours evolve more quickly than cuckoos host-switch. Evidence suggests that cuckoos host-switch on the order of decades, but that eggshell colours generally^56^ change more slowly^57^ likely due to the relatively low heritability of egg colouration^58^. In fact, sex-linked traits such as the cuckoo’s eggshell colour^12^ and female cuckoo’s polymorphic plumage colour^59^, are expected to change even more slowly^60^.

These dynamics are also complex from the hosts’ perspective, which ultimately drives the evolution of egg colour mimicry. For example, while most hosts use eggshell colour as an egg rejection cue, their rejection does not always correspond to the perceived colour differences between their own and experimentally introduced eggs^20,21,24^. Instead, hosts generally respond more strongly to eggs browner than their own^20,61^. Naturally, such behaviours should impact how cuckoo colour mimicry evolves and would result in some colours being more mimicable than others. While egg recognition is exceptionally well-studied^9^, there are still substantial gaps in our understanding of egg recognition. For example, background adaptation^62^ can make it more challenging to discriminate two bluish eggs on a brown background than two brown eggs. Unfortunately, current perceptual models do not account for these properties^62–65^, nor the fact that discrimination thresholds vary over the avian colour space^66,67^ in as-of-yet unknown ways^68,69^.

Finally, studies that have tracked banding data imply that cuckoos disperse relatively far from their natal ranges^70^, ultimately exporting their genes to new regions. Thus, even if young female cuckoos parasitize their natal host species later in life, they may face new host populations that may have different eggshell traits^71,72^ or recognition abilities^31,73^ than their natal host population. Generally, we assume host populations share the same discrimination threshold (i.e., rejecting cuckoo eggs that differ from host egg colours by a certain number of just noticeable differences); however, this is not a valid assumption because host populations can differ significantly in their recognition abilities^31,73,74^ and discriminable differences depend upon the specific colours being compared^68,69^. Although not yet fully understood^68^, these factors should negatively impact a cuckoo lineage’s ability to adapt egg mimicry and may impact the evolution of egg recognition in a number of ways^68^.

### Coevolutionary alternation

Despite cuckoo-host dynamics fueling our understanding of coevolution for decades^2,3,15,75,76^, few studies have explored whether alternative coevolutionary processes can explain the apparent mimicry between cuckoo and host eggs^34^. Coevolutionary alternation is one such coevolutionary process^77^. Under this coevolutionary process, cuckoo lineages engage in a range of short-term coevolutionary relationships with a range of hosts until they evolve sufficiently competent egg recognition abilities. Then, cuckoos switch to new, potentially naïve, hosts (Fig. 3). Cuckoo eggshell features would be selected at each iteration of this process, although their eggs may already be similar in colour to their new hosts’ eggshell colours. Likewise, their new hosts may or may not have evolved egg recognition from prior experiences with other cuckoo lineages. Thus, cuckoo eggshell features are impacted by (often unknown) historic host species in a variable process of coevolutionary lag^40,78,79^. This would result in cuckoos with egg colours that provide a generalized match to many potential hosts, resulting in less specialized traits than under coevolutionary arms races^80,81^. Therefore, we expect a high rate phenotypic divergence (i.e., many cuckoo egg colours), but not a high rate of host-race divergence, a pattern which was recently supported^29,but^ ^see, 76^. Resultantly, cuckoos produce just a few variable egg types that get recycled among their current and potential host species. Determining which coevolutionary process governs cuckoo-host dynamics, is challenging given that it depends on largely unknown, historic, events^7^.

**Figure 3.**
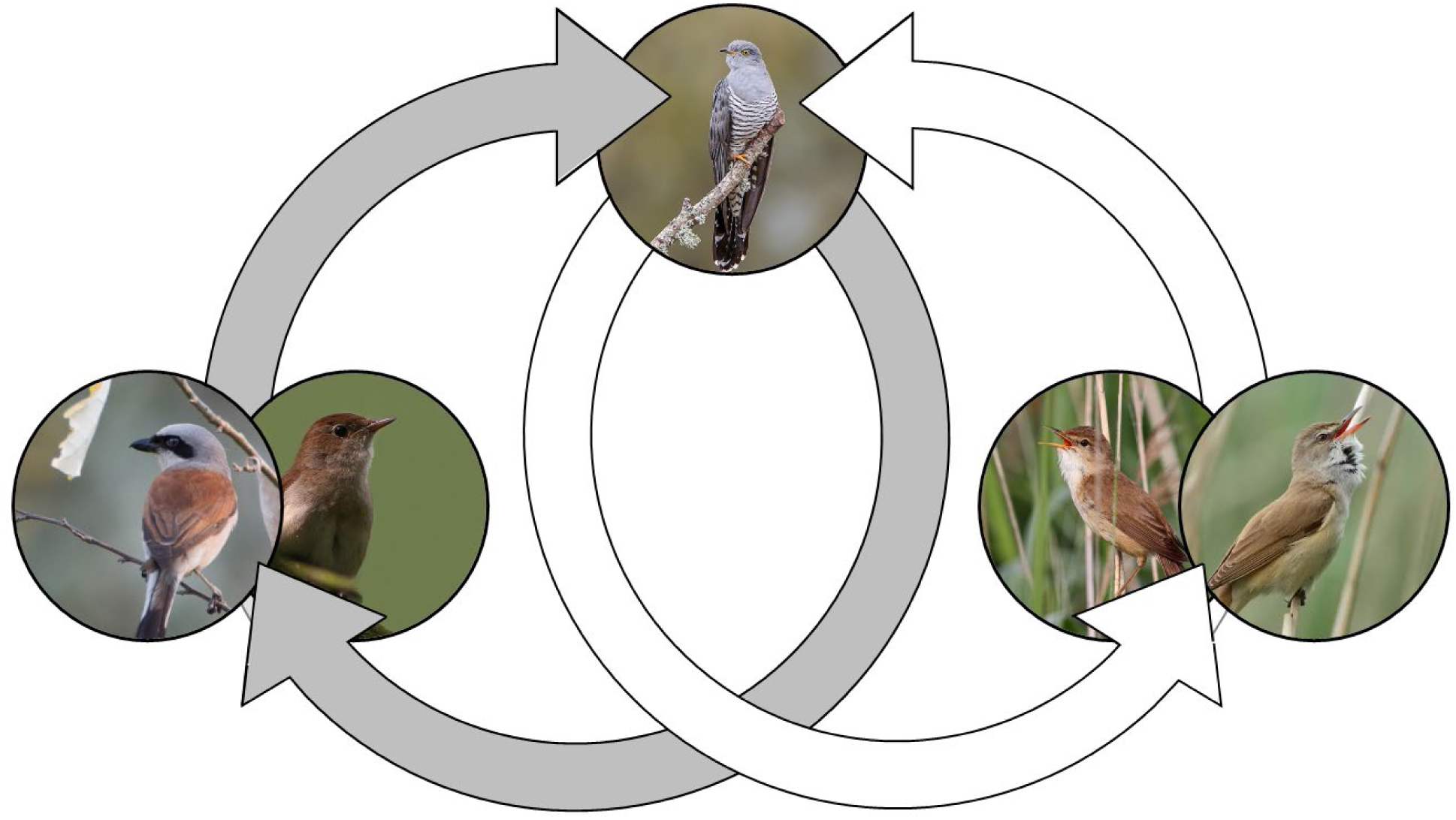
Coevolutionary alternation hypothesis for egg colour mimicry in Common cuckoo. Due to a continuous cycle of host-switching, cuckoos exploit an ever-changing community. As once-common hosts evolve strong egg recognition (grey arrows), cuckoos switch to alternative host species (white arrows). Cuckoo lineages carry eggshell phenotypes they have evolved from their varied historic hosts to new hosts. Ultimately, these evolutionary admixtures select for a generalized eggshell phenotype that generally works well, but is most similar to only a subset of their host community. Photo credits: <u>Cuckoo</u> by Andy Morffew (<u>CC-BY-2.0</u>), <u>Reed warbler</u> by Frans Vandewalle (<u>CC-BY-NC-2.0</u>), <u>Great reed warbler</u> by Peter Rohrbeck (<u>CC-BY-SA-4.0</u>), <u>Nightingale</u> by Petra Karstedt (<u>CC-BY-2.0</u>), and <u>Red-backed shrike</u> (public domain, <u>CC0 1.0</u>).

While broadly similar, coevolutionary arms races and coevolutionary alternation would generate substantial differences in cuckoo egg morphology, and in turn the ability of cuckoo host-races to specifically mimic the eggshell colours of their hosts’ eggs. Coevolutionary alternation implies a process of continual host-switching that will select for less specialized cuckoo eggs than under coevolutionary arms races^77^. While we expect that the cuckoo eggs will still have relatively diverse colouration^29^, they will generally share similar colours regardless of their host species. As a result, under coevolutionary alternation mimicry will directly relate to host egg colour, such that the cuckoo’s eggshell colours will match their hosts when their hosts lay eggs of these generalized eggshell colours, but they will not match more colourful host eggs^33^. Moreover, because cuckoo host-races can result from various recent or ancient host-switching events^31^, under this model of coevolution cuckoo host-races should not necessarily produce distinct eggshell colours^39^. In fact, coevolutionary alternation implies that cuckoos will lay a more generalized egg phenotype that may not specifically match their current hosts’ eggs. Instead, cuckoos should often match the egg colours of other (potentially former) hosts just as well as their current host.

## Conclusion

We have long assumed that the perceived similarity between host and cuckoo eggs was evidence of independent coevolutionary arms races; however, similarity between host and cuckoo eggshell colours does not necessarily imply any coevolved mimicry^82,83^ and there are other coevolutionary processes besides arms races that can give rise to these same patterns^84–87^. Here, we explore host and cuckoo eggshell colouration to test several predictions of coevolutionary arms races using an expansive dataset of eggshell reflectance spectra from parasitized host clutches. Specifically, we found that because cuckoo eggs did not mimic extreme host eggshell colours, the degree of mimicry was directly (and strikingly) related to host eggshell colouration (Fig. 1). Furthermore, only three of 39 cuckoo host-races were more likely to match their specific hosts’ eggshell colour than the eggshell colour of other host species. Of these, the only common host-race to specifically match its host was the cuckoo host race parasitizing the European robin. Similarly, only two of the 39 cuckoo host-races laid eggs that had colours distinct from other cuckoo host-races. In this case, the cuckoo parasitizing the redstart was the only common host-race to lay an egg distinct from those laid by other host-races. Thus, across these 39 host species, we did not find evidence of coevolutionary arms races, in either our newly collected data, a classic dataset, or the combined dataset.

Collectively, our data support the alternative coevolutionary alternation hypothesis which states that cuckoo eggshell colours evolve through a pattern of continuous host-switching. This coevolutionary process is logical because cuckoos frequently switch hosts^12^ and maternally derived eggshell colours are likely slow to evolve. Ultimately, this process gives rise to a more generalized egg phenotype that sometimes matches the phenotypes of particular hosts well but is unlikely to evolve highly specialized eggshell phenotypes that mimic their particular hosts. In evolutionary biology we often classify coevolutionary relationships (e.g., cuckoo-host, pollinator-plant relationships) along a continuum of specialization ranging from highly specialized to more generalized. Coevolutionary arms races selects for highly specialized phenotypes, while coevolutionary alternation selects for far more generalized phenotypes. While our findings challenge the accepted paradigm of a classic system, we argue that in doing so our findings uncover important and under-appreciated information about the process of coevolution. It is clear that cuckoos and their hosts will continue to teach us much about the patterns, pace, and process of coevolution, which will undoubtedly continue to fuel our understanding of evolution for decades to come.

## Methods

### Eggshell reflectance measurements

We collected eggshell reflectance for 38 host species and their corresponding cuckoos (N = 732 host eggs from 220 parasitized clutches) at two natural history museums: Western Foundation of Vertebrate Zoology, and Delaware Museum of Nature and Science. We acknowledge that eggshell colour can degrade over time in museum collections, with gradual decreases in brightness reflectance and chroma^88^; however, because we measured both host and cuckoo eggs from the same parasitized clutches, both eggs experienced identical storage conditions and duration. This paired design ensures that any degradation affects both eggs similarly, preserving the relative colour differences that are central to our mimicry assessments. In addition, for illustrative purposes (to localise the ‘colour’ of a white egg), we collected the spectral reflectance of two freshly collected mourning dove eggs, which were used to determine the ‘colour’ of a pure white egg. These reflectance spectra were collected using a portable spectrophotometer (Jaz-PX, Ocean Optics) with a built in pulsed xenon light source (Jaz-PX, Ocean Optics) and a 600µm bifurcating fibre optic. The fibre optic terminated in a probe tip over which we placed a customized probe tip. Using this system, we recorded egg reflectance spectra at a consistent 45° coincident oblique measurement angle^89^. We calibrated the spectra to a fresh white diffuse standard of 99% reflectance (WS-1-SL, Ocean Optics) and a customized dark reference. Then for each egg we produced an average spectrum from six measurements across the eggshell surface avoiding spots whenever possible: twice at the blunt end, twice at the equator, and twice at the sharp end^89^. These data were supplemented with eggshell reflectance spectra from a previously published study^4^, that were provided courtesy of the authors. This dataset contained one spectral reflectance measurement as a representative of each host clutch (N = 249) and a spectrum of the cuckoo egg found in that clutch (N = 249). Together, these reflectance spectra represent the egg colours of 39 host species, from 468 parasitized clutches including 6 doubly parasitized clutches (N = 980 host eggs, N = 474 cuckoo eggs; Table S4). Thus, our dataset includes common and rare hosts. All reflectance spectra were truncated and interpolated to evenly span the avian visible range of 300 to 700 nm.

### Visual models and colour spaces

For these analyses, we used the common blackbird *Turdus merula* as general representative of a typical ultraviolet sensitive viewer. We estimated spectral sensitivity curves^90^ using data on their peak sensitivity, oil droplet transmission filter cut-offs, and ocular media transmission, which were derived from microspectrophotometry^91^. Then, to estimate avian perceived colour we calculated the quantum catches for each spectral reflectance (host eggs, cuckoo eggs, all birds’ eggs, and for the colour library) using the following equation

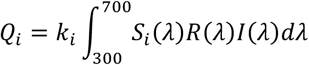

Where, where *Si*(λ) is the normalized spectral sensitivity of photoreceptor *i*, *R*(λ) is the stimulus reflectance, *I*(λ) is the irradiance spectrum. In this case, we used a radiometrically calibrated radiometer (StellarRad, Stellarnet) to measure irradiance above the nests of nine open cup nesting species and used the average irradiance spectrum as a representative irradiance spectrum. For these analyses, we used a scaling coefficient, *k_i_*, to apply a von Kries correction, which is generally believed to approximate the colour perception of animals with colour constancy^92^.

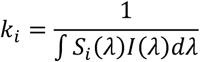

This resulted in similar quantum catch estimates as using an idealized irradiance (set to 1 over all wavelengths), which is expected when using a von Kries correction^93^.

Here, we depict eggshell colouration within an avian visual colour space, known as the tetrahedral colour space^93^. This space represents the relative stimulation of each of the avian-viewer’s four photoreceptors. Birds’ eggs fall on a line within this colour space because most of the variation in colours is due to two pigments^26^: biliverdin that generates blue-green and protoporphyrin that generates brown. In this case, most (75%) of the variation colouration was explained by the x-coordinate within this colour space across all host and cuckoo eggs (N = 1,454). Thus, in this case, we will use this as a metric to represent eggshell colour.

Finally, we used the receptor noise limited (RNL) model^64^ for all colour comparisons. This model calculates how different the colours of two stimuli (in this case, eggs) would appear to hosts. We set the Weber fraction at 0.10 for the long-wavelength sensitive photoreceptor, based on behavioural experiments^94^ and used data on the blackbird’s relative photoreceptor abundance^91^. These models report perceived differences in units of just noticeable difference (JND), where a value of 1 JND is the threshold beyond which colours are perceivably different. Larger JND values represent larger perceived differences, while those less than 1 are not perceivably different. We used these models to calculate the perceived eggshell colour mimicry between host eggs and the specific cuckoo eggs found in their nests; however, in addition to comparing cuckoo eggs to host eggs, we also used RNL models to compare cuckoo eggs to other cuckoo eggs (comparisons between host-races) and to compare host eggs to other host eggs (interspecific comparisons).

### Statistical analyses

#### Quantifying variation in eggshell colour

We quantified the diversity of eggshell colours within the tetrahedral colour space^93^. If cuckoos’ eggshell colours have evolved via coevolutionary arms races, we expect that their eggshell colours would be nearly as diverse as those of their hosts. By contrast, we may find that cuckoos only match a few host egg colours suggesting that arms races may not operate for all of their hosts. Although colour volumes are often reported as the volume of the convex hulls encompassing a species’ coordinates within the colour space^56,95,96^, this approach is inappropriate in our case because the volume is determined solely by rare colours and increases monotonically with the number of points^97^. Instead, we summarized the variability of coordinates for cuckoo and host eggshell colours within the tetrahedral colour space using covariance matrices, the determinants of which scale proportionally with colour volumes^98^ and are directly comparable between cuckoo and host colours. Thus, larger determinants represent clouds of colour coordinates with larger volumes, while smaller determinants represent egg colours that are less variable. Here we used a bootstrap approach to statistically compare relative colour volumes for hosts’ eggs and cuckoo eggs (overall, and individual species comparisons).

#### Is mimicry related to host eggshell colour

To test whether cuckoo eggshell colour mimicry was predicted by the hosts’ eggshell colours (the x-coordinate of the tetrahedral colour space), we used a linear mixed model (LMM) fitted via Markov Chain Monte Carlo using the R package MCMCglmm^99^. Our model accounts for phylogenetic non-independence and for dependencies within species and clutches, which allowed us to examine data from all clutches. In this case, we acquired a set of 100 phylogenetic hypotheses representing the relationships among all host species from birdtree.org^100^. These phylogenetic hypotheses were summarized to a single maximum clade credibility tree, and we then constructed the phylogenetic covariance matrix from this tree. Our LMM predicted eggshell colour mimicry by eggshell colour and squared eggshell colour (i.e., a second order polynomial regression), as well as the source of the data (‘NHM’, ‘DMNH’, or ‘WFVZ’). This model included species and clutch ID as random effects. We ran an effective number of 5,000 MCMC iterations (55,000 iterations in total, discarding the first 5,000 for burn-in and retaining every 10^th^ iteration). In addition to this larger global analysis, we ran separate models using only our newly collected dataset and only the classic data^4^. These models were similarly constructed, except the model for data from the classic dataset^4^ did not require a random effect for the clutch since their study included only one host egg per clutch. By contrast, in the global analysis we accounted for the dependency between host eggs from the same clutch.

#### Within and between host and host cuckoo host-race comparisons

We further evaluated how adapted cuckoo eggshell colours are to their host race’s eggshell colours. We first derived the perceived difference between all cuckoo eggshells, across all clutches, cuckoo-host races and studies. We then fitted cuckoo-host race specific LMMs to quantify the average differences between cuckoo eggshells from that specific cuckoo-host race (average “within” difference) and between cuckoo eggshells from this cuckoo-host race to all other cuckoo-host races (average “between” difference). The LMMs included random effects for the other cuckoo-host race, the clutch ID and the source of the data (‘NHM’, ‘DMNH’, or ‘WFVZ’). The random effects were set up to account for dependencies among all four eggs contributing to each pair of distances. We fitted the models using MCMC with an effective number of 5,000 MCMC samples, from which 95% Bayesian highest posterior density (HPD) intervals for the average within and between differences were derived. In addition to these models for the perceived differences between cuckoo eggshells, we fitted similar models to the perceived differences between host eggshells, as well as to the perceived differences between cuckoo and host eggshells.

## Data availability

All the data will be made available during peer-review and publicly thereafter.

## Acknowledgements

We thank MC Stoddard and M Stevens for providing access to original source data, and ME Hauber and K Szala for insightful comments on an earlier draft. In addition, we are grateful to the Western Foundation of Vertebrate Zoology, and the Delaware Museum of Nature and Science for access to their collection and to L Hall, R Corado, and M Halley for facilitating our collection visits.

## Author contributions

Conceived of the study: DH; developed methods: DH; collected data: DH and JV; analysed the data: DH, DK; wrote the paper: DH, AN, with help from JVAV and JV; final editing: all authors.

## Additional information

Electronic supplementary materials Supplemental figures

---

**Figure S1.**
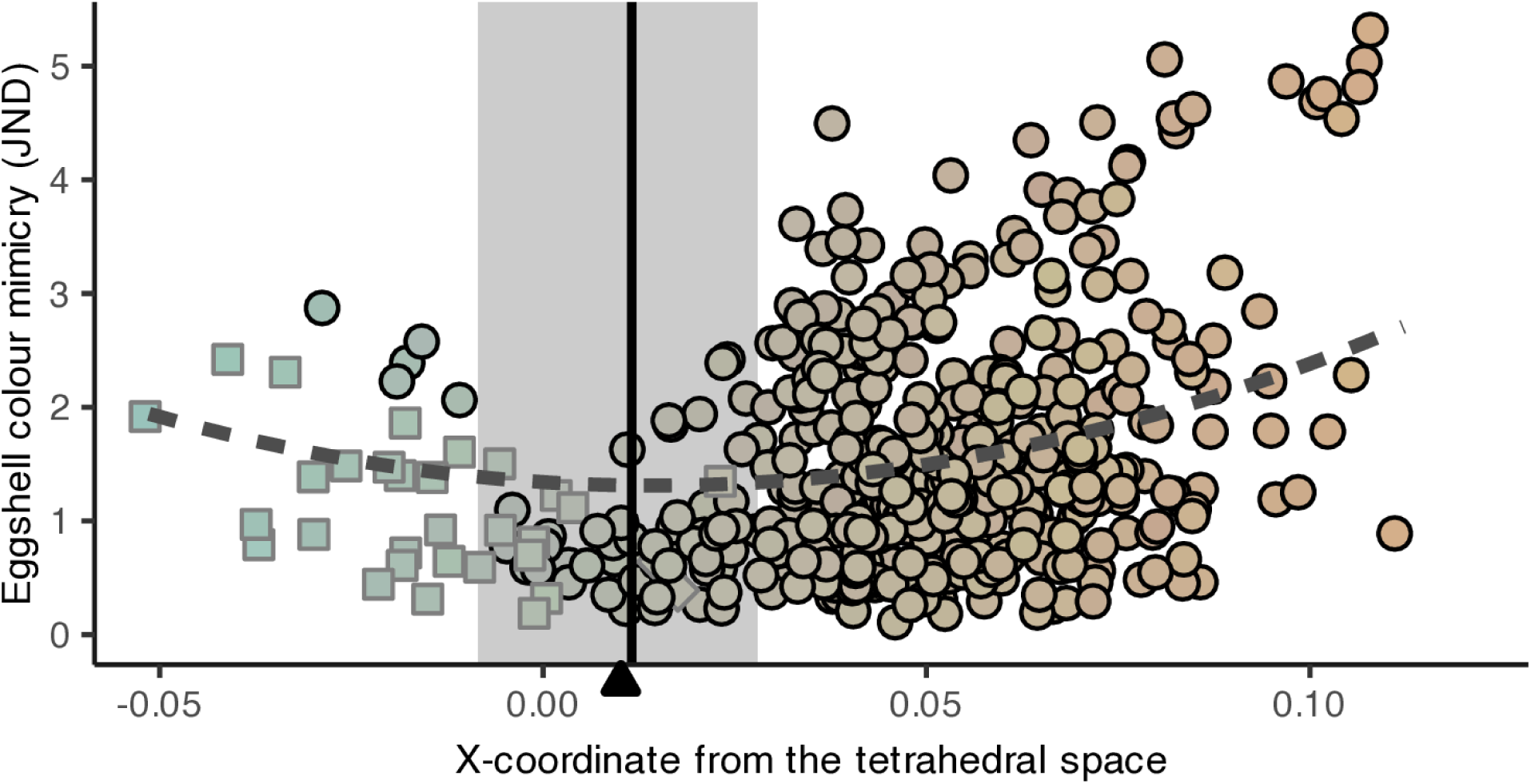
The mimicry of cuckoo eggshell colours to their specific hosts compared to colours of each cuckoo eggshell. Here we illustrate the nonlinear fit and plot all points in the colours of the cuckoo’s eggs and highlight the best matched cuckoo eggshell colour (vertical solid line). The eggshell colour of a pure white mourning dove egg (filled triangle) is very close to this best match and within the 95% confidence interval (grey area) estimated from the linear mixed effect model.

**Figure S2.**
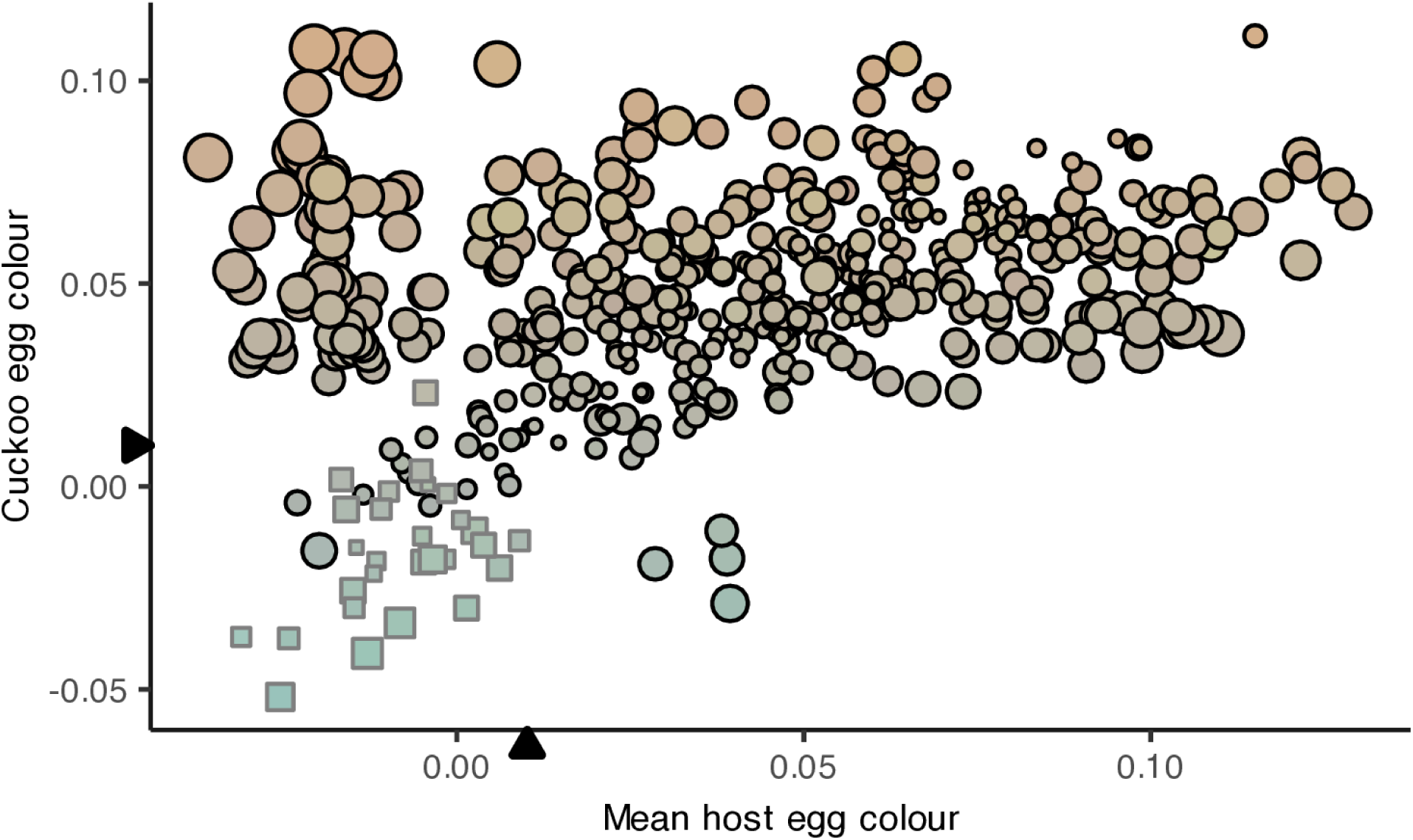
A comparison between host and cuckoo eggshell colour. Here we plot the colour, defined as the x-coordinate from the tetrahedral colour space, for both hosts and their respective cuckoos. Host eggshell colours are calculated as the average colour within each host’s clutch, while cuckoo eggshell colours represent the colour of the cuckoo egg(s) from each host clutch. To aid interpretation all points are depicted in cuckoo eggshell colours. We show redstarts and the redstart-type cuckoo host-race as squares with grey outlines, because this was the only common host-race that had an egg with a colour distinct from those laid by other cuckoo host-races (Fig. 2). The coordinate of a mourning dove egg is shown (filled triangles) along both axes to indicate the coordinate associated with a pure white eggshell. Finally, all points are size scaled according to the achieved eggshell mimicry within the clutch, such that smaller points represent better colour matches between the host clutches and the specific cuckoo that parasitized them.

**Figure S3.**
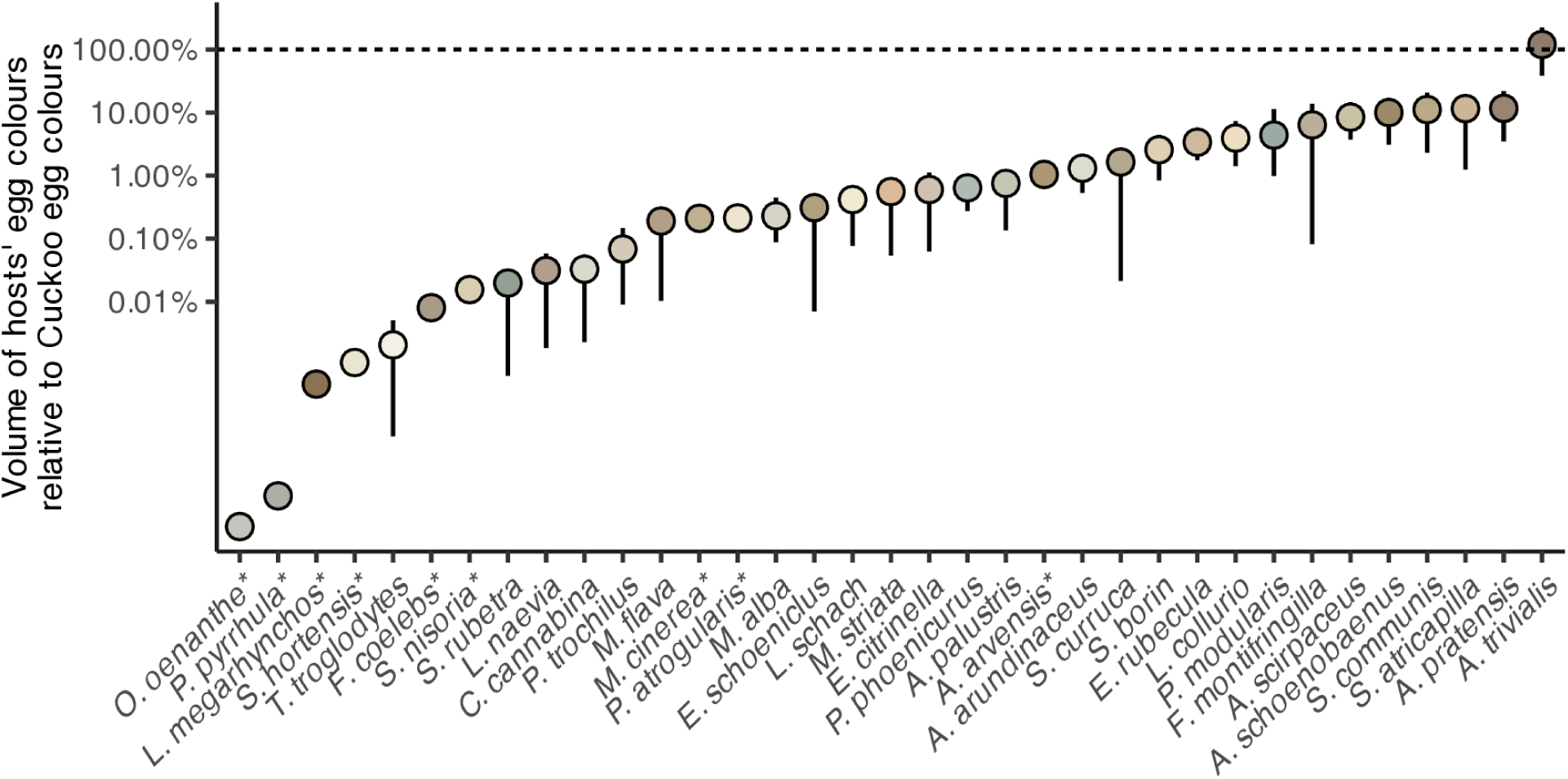
Differences in volumes of hosts’ egg colours to cuckoo egg colours (pooling all host-races), as the ratio of the determinants of the respective covariance matrices of colours in the tetrahedral space. The vertical axis is shown on the log-scale. For species marked by an asterisk (*), too few (< 10) eggs were available to estimate confidence intervals by bootstrap. Four rare host species with three or less eggs were excluded because the covariance matrix in the tetrahedral space could not be reliably estimated (see Fig. S4 and Table S2).

**Figure S4.**
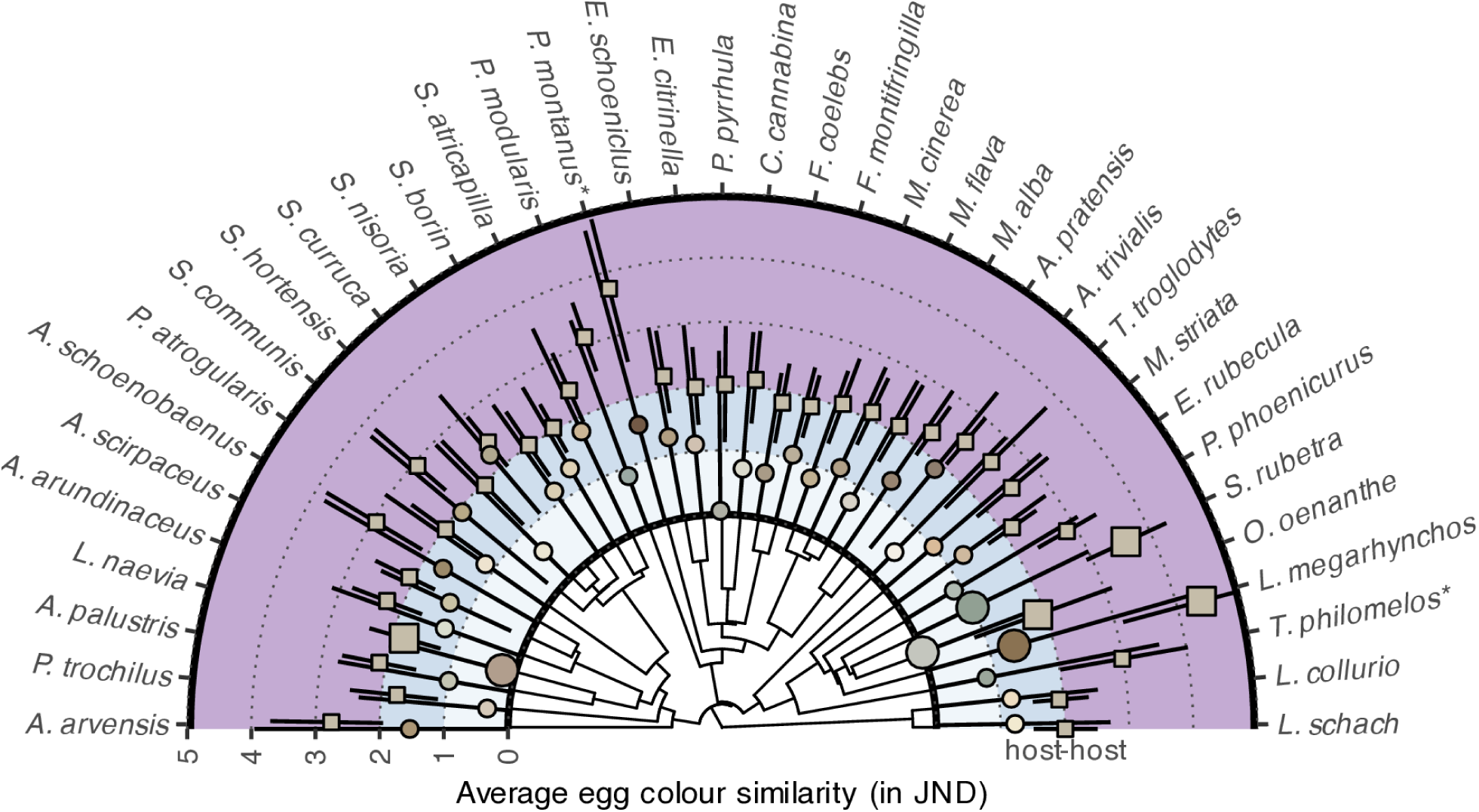
Average perceived difference in eggshell colour (in JND) among 37 host species. Here, we depict mean perceived colour differences (with 95% confidence intervals) among host species along the edges of a maximum clade credibility tree. Egg colour differences within host species are shown in circles, while differences between host species are shown in squares. Enlarged points represent significant differences. The bands of colour along the outer ring represent differences < 1 JND (white), ≤ 2 JND (pale blue), and > 2 JND (light purple). The circles are coloured based on the average eggshell colour for each host species, while the colours of the squares represent the average colour across all host species. Asterisks indicate hosts with sparse sampling that should be interpreted with care. Seven rare host species were excluded because there were too few specimens to reliably estimate within host variation (Table S2).

**Figure S5.**
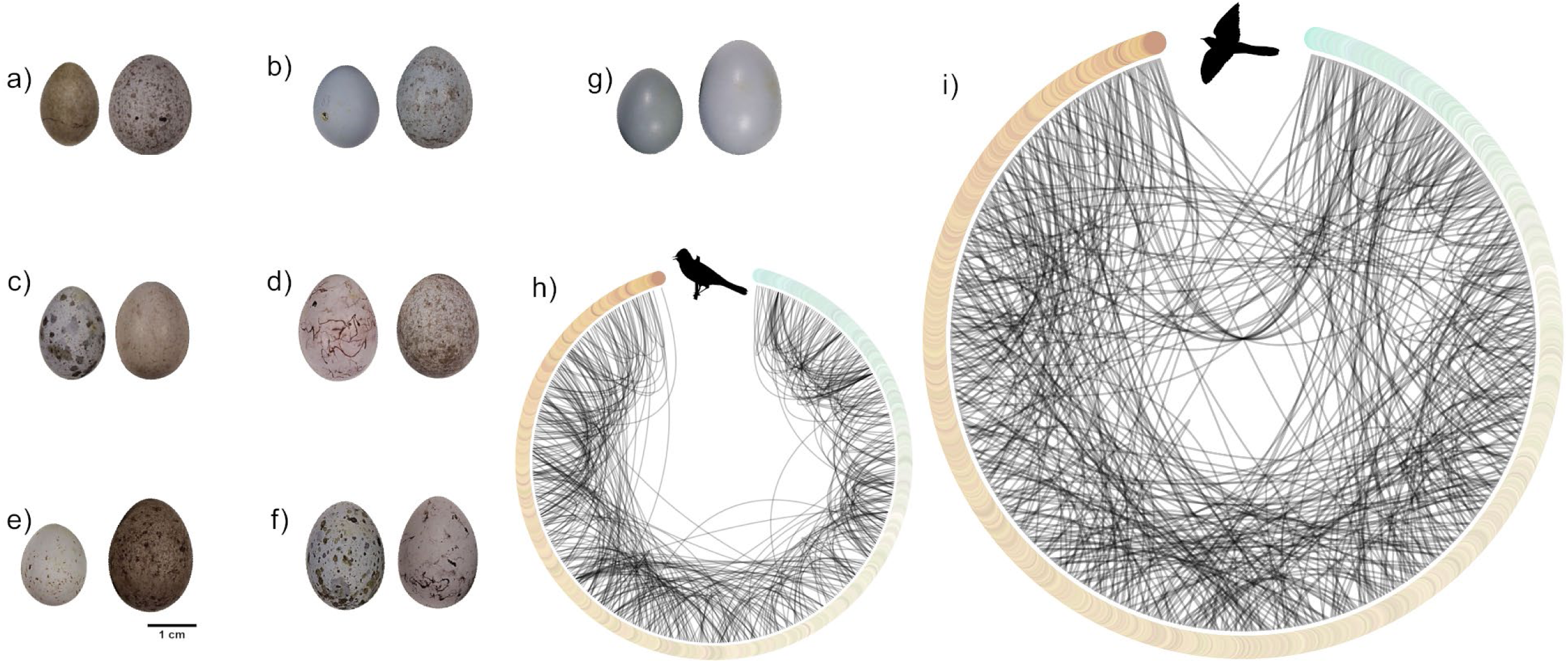
Failures to achieve mimicry are common across all of the cuckoo’s hosts. Here we illustrate the matches between host (left) and cuckoo (right) eggs from the a) sedge warbler *Acrocephalus schoenobaenus* (wfvz 23347), b) dunnock *Prunella modularis* (wfvz 151020), c) marsh warbler *Acrocephalus palustris* (wfvz 23386), d) yellow bunting *Emberiza citrinella* (wfvz 23161), e) linnet *Linaria cannabina* (wfvz 151019), f) great reed warbler *Acrocephalus arundinaceus* (wfvz 25202), and g) whinchat *Saxicola rubetra* (wfvz 23443). Photographs (a-g) obtained through the collection database for the Western Foundation of Vertebrate Zoology (https://collections.wfvz.org/). Eggshell colours vary from blue-green to brown, which we illustrate in h) a circular plot for hierarchical edge bundling. The edges represent the mean eggshell colours of 468 host clutches, while the lines connect each host clutch to another host clutch from the same species that is closest to the median colour contrast (JND) to all other clutches. The arc represents the chromatic contrast (JND) between any pair of clutches, such that larger colour differences will pass through the centre of the plot. Since eggshell colours are broadly similar most linkages are local and do not span the plot. By contrast, i) cuckoo eggshell colours are not closely associated with the colours of the clutches that they parasitize. Here, we illustrate 474 cuckoo eggs are arranged by their eggshell colours alongside the 468 host clutches (6 clutches were parasitized by two cuckoos), connecting each cuckoo egg to the specific host that it parasitized. It is clearly visible that cuckoos rarely match their hosts’ eggshell colour. Inset icons of hosts represented by the silhouette of a *Sylvia atricapilla* (credit: Alexis Lour) and a common cuckoo (credit: Chris Romeiks) downloaded at https://www.phylopic.org/. Inset egg photographs were taken by René Corado and used with permission (photo copyright: René Corado and Western Foundation of Vertebrate Zoology, all rights reserved)

**Table S1.**
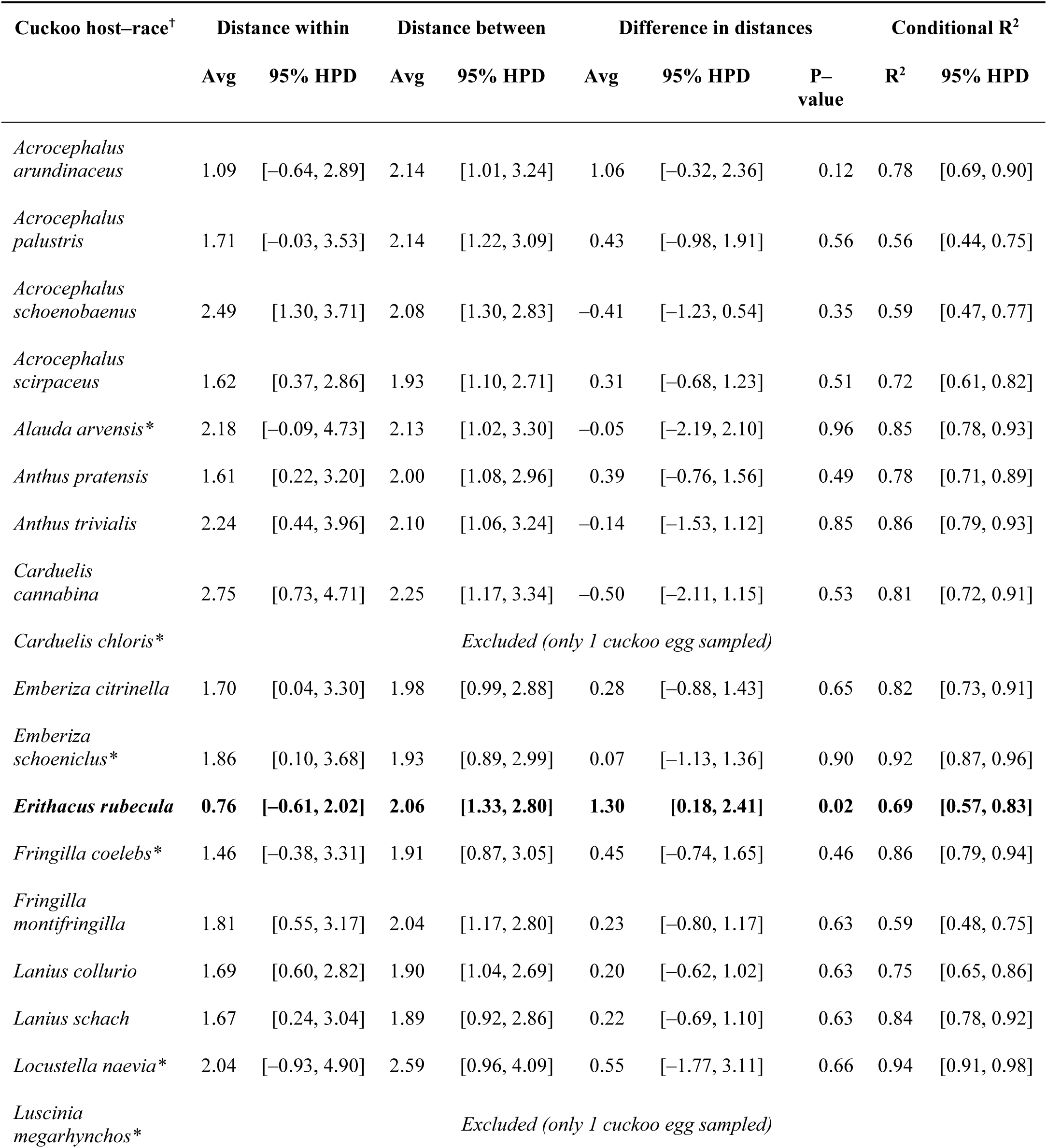
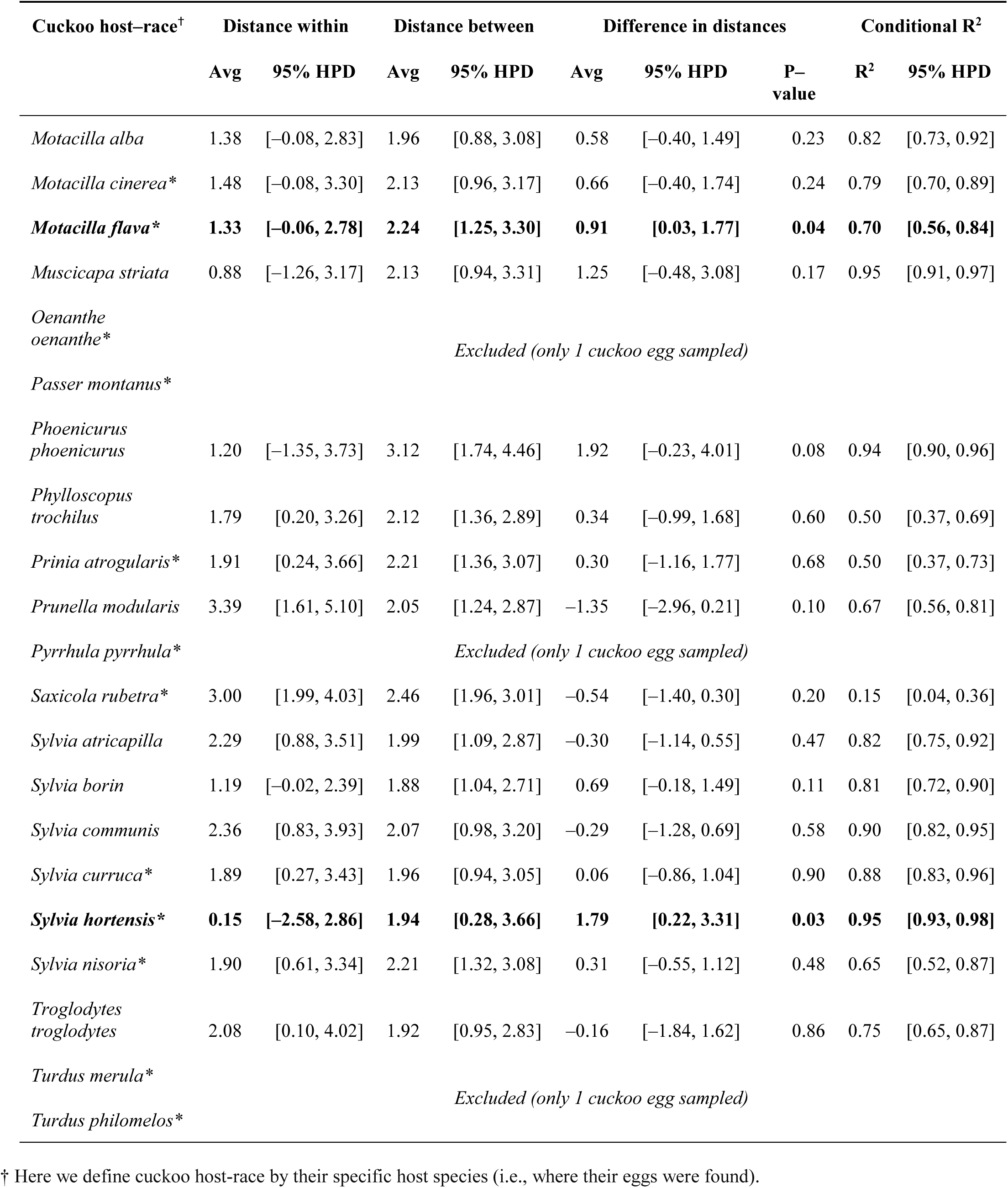
Average colour distances (in JND) between cuckoo eggs and their specific hosts’ eggs (within), between cuckoo eggs and the eggs of all other hosts (between), and the average difference in these distances. Average values are accompanied by 95% highest posterior density (HPD) intervals. Host–races with significant differences at the 5% level are highlighted in bold. Host–races with 3 or less eggs are marked by an asterisk (*).

**Table S2.**
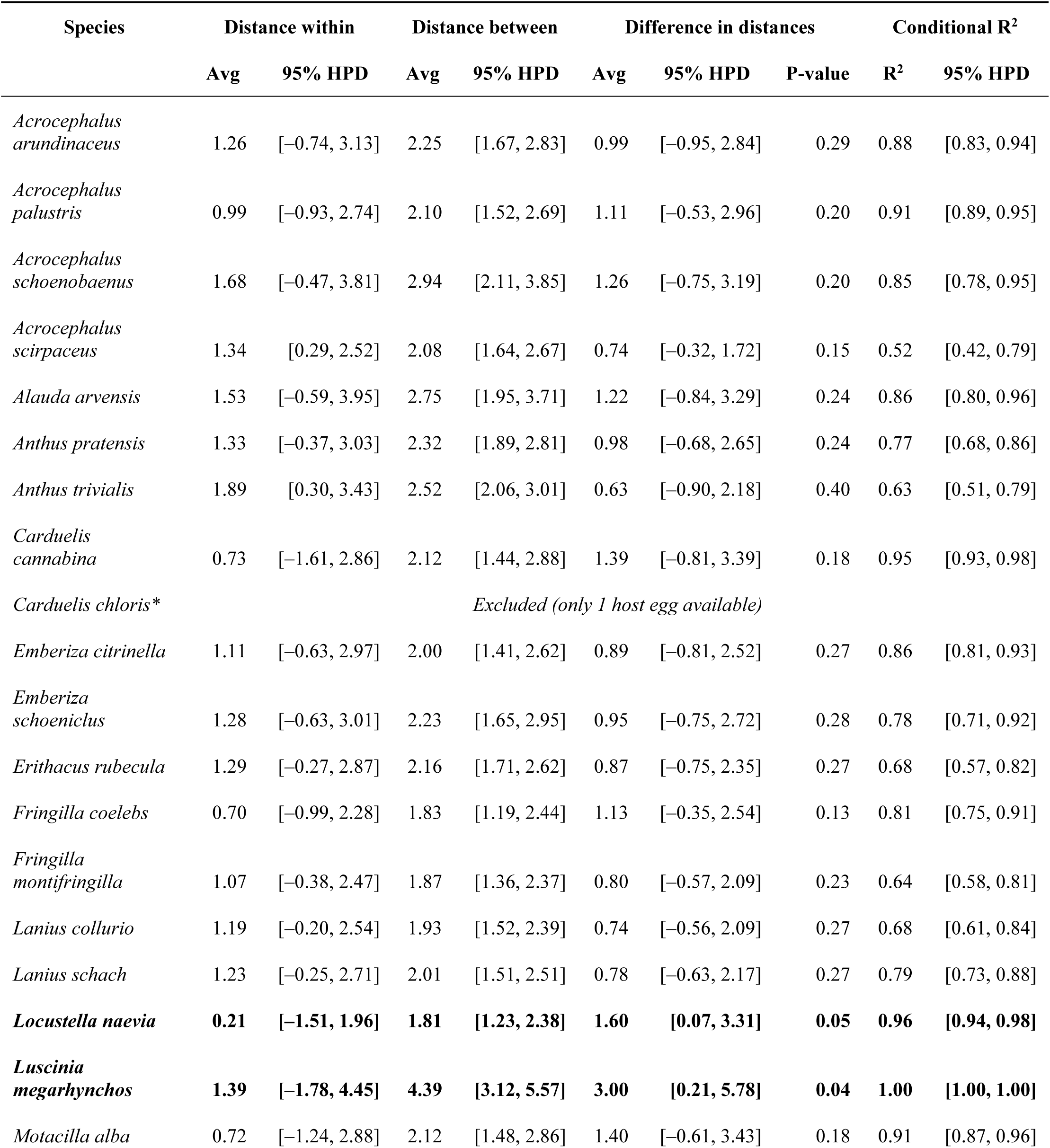
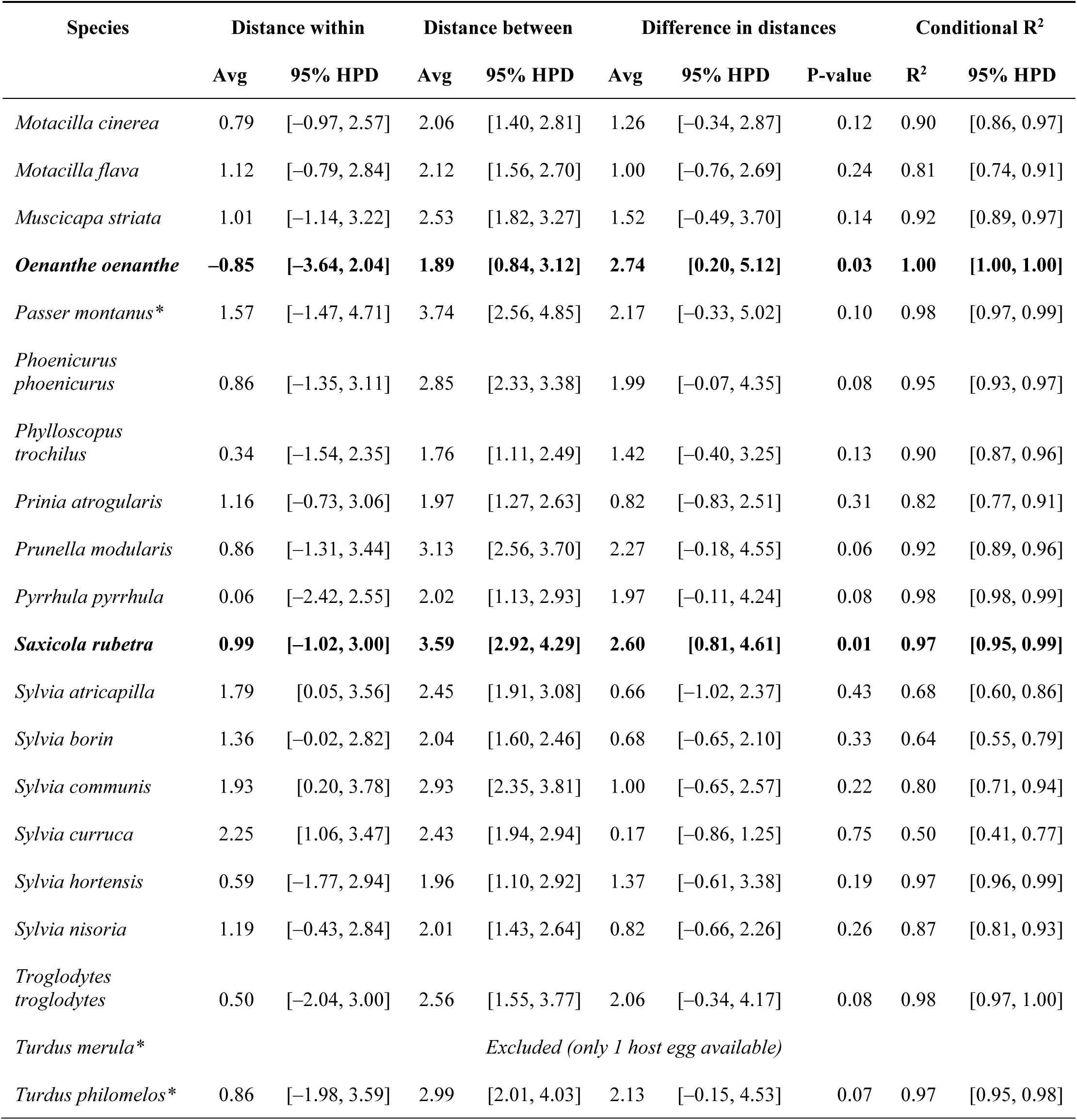
Average colour distances (in JND) between hosts’ eggs within the same species (within), between hosts’ eggs from different species (between), and the average difference in these distances. Average values are accompanied by 95% highest posterior density (HPD) intervals. Species with significant differences at the 5% level are highlighted in bold. Species with 5 or less eggs are marked by an asterisk (*).

**Table S3.** Average colour distances (in JND) between cuckoo eggs within the same host–race (within), between cuckoos of different host–races (between), and the average difference in these distances. Average values are accompanied by 95% highest posterior density (HPD) intervals. Host–races with significant differences at the 5% level are highlighted in bold. Host–races with 3 or less eggs are marked by an asterisk (*).

| Cuckoo host–race<sup>†</sup> | Distance within |  | Distance between |  | Difference in distances |  |  | Conditional R<sup>2</sup> |  |
| --- | --- | --- | --- | --- | --- | --- | --- | --- | --- |
|  | Avg | 95% HPD | Avg | 95% HPD | Avg | 95% HPD | P-value | R<sup>2</sup> | 95% HPD |
| *Acrocephalus arundinaceus* | 1.56 | [0.73, 2.40] | 1.93 | [1.47, 2.56] | 0.37 | [−0.21, 1.01] | 0.22 | 0.57 | [0.50, 0.87] |
| *Acrocephalus palustris* | 1.69 | [0.85, 2.48] | 1.78 | [1.38, 2.18] | 0.09 | [−0.58, 0.72] | 0.78 | 0.25 | [0.18, 0.58] |
| *Acrocephalus schoenobaenus* | 1.71 | [0.70, 2.68] | 1.60 | [1.17, 2.07] | −0.11 | [−0.94, 0.78] | 0.79 | 0.51 | [0.41, 0.79] |
| *Acrocephalus scirpaceus* | 1.02 | [0.01, 1.89] | 1.33 | [0.95, 1.73] | 0.31 | [−0.56, 1.17] | 0.46 | 0.63 | [0.53, 0.82] |
| *Alauda arvensis* * | 1.01 | [−0.66, 2.64] | 1.30 | [0.47, 2.15] | 0.29 | [−0.96, 1.63] | 0.64 | 0.78 | [0.69, 0.96] |
| *Anthus pratensis* | 1.01 | [−0.04, 2.08] | 1.45 | [1.05, 1.85] | 0.45 | [−0.59, 1.40] | 0.36 | 0.69 | [0.62, 0.84] |
| *Anthus trivialis* | 1.33 | [−0.07, 2.92] | 1.52 | [0.73, 2.51] | 0.19 | [−0.86, 1.37] | 0.73 | 0.83 | [0.78, 0.98] |
| *Carduelis cannabina* | 1.48 | [0.01, 3.10] | 1.49 | [0.64, 2.44] | 0.01 | [−1.18, 1.16] | 0.99 | 0.76 | [0.67, 0.96] |
| *Carduelis chloris* * | Excluded (only 1 cuckoo egg sampled) |  |  |  |  |  |  |  |  |
| *Emberiza citrinella* | 0.94 | [−0.46, 2.17] | 1.23 | [0.61, 1.90] | 0.29 | [−0.70, 1.36] | 0.58 | 0.73 | [0.63, 0.92] |
| *Emberiza schoeniclus* * | 0.52 | [−1.05, 2.19] | 1.14 | [0.33, 2.01] | 0.62 | [−0.51, 1.78] | 0.28 | 0.92 | [0.88, 0.98] |
| *Erithacus rubecula* | 1.35 | [0.36, 2.33] | 1.49 | [1.06, 1.92] | 0.14 | [−0.77, 1.03] | 0.74 | 0.58 | [0.48, 0.83] |
| *Fringilla coelebs* * | 1.31 | [−0.51, 3.16] | 1.37 | [0.46, 2.40] | 0.06 | [−1.26, 1.35] | 0.93 | 0.79 | [0.70, 0.97] |
| *Fringilla montifringilla* | 1.46 | [0.69, 2.20] | 1.62 | [1.23, 2.00] | 0.17 | [−0.40, 0.77] | 0.56 | 0.43 | [0.36, 0.69] |
| *Lanius collurio* | 1.01 | [0.11, 1.91] | 1.36 | [1.00, 1.70] | 0.35 | [−0.42, 1.22] | 0.40 | 0.63 | [0.55, 0.81] |
| *Lanius schach* | 0.87 | [−0.11, 1.98] | 1.47 | [0.95, 1.95] | 0.59 | [−0.17, 1.31] | 0.11 | 0.73 | [0.68, 0.85] |
| *Locustella naevia* * | 1.88 | [−0.22, 4.16] | 2.20 | [0.91, 3.68] | 0.32 | [−1.16, 1.71] | 0.65 | 0.93 | [0.90, 1.00] |
| *Luscinia megarhynchos* * | Excluded (only 1 cuckoo egg sampled) |  |  |  |  |  |  |  |  |
| *Motacilla alba* | 1.29 | [0.40, 2.27] | 1.72 | [1.19, 2.23] | 0.42 | [−0.32, 1.22] | 0.27 | 0.67 | [0.61, 0.90] |
|  | Avg | 95% HPD | Avg | 95% HPD | Avg | 95% HPD | P-value | R<sup>2</sup> | 95% HPD |
| *Motacilla cinerea* * | 1.64 | [0.01, 3.26] | 1.44 | [0.71, 2.20] | −0.20 | [−1.57, 1.24] | 0.79 | 0.62 | [0.48, 0.93] |
| *Motacilla flava* * | 1.59 | [0.12, 3.02] | 1.61 | [0.95, 2.44] | 0.02 | [−1.18, 1.27] | 0.98 | 0.50 | [0.37, 0.89] |
| *Muscicapa striata* | 0.44 | [−1.26, 2.29] | 1.21 | [0.24, 2.36] | 0.77 | [−0.54, 2.05] | 0.23 | 0.94 | [0.92, 0.99] |
| *Oenanthe oenanthe* * | Excluded (only 1 cuckoo egg sampled) |  |  |  |  |  |  |  |  |
| *Passer montanus* * |  |  |  |  |  |  |  |  |  |
| ***Phoenicurus phoenicurus*** | **1.49** | **[0.62, 2.44]** | **3.31** | **[2.85, 3.79]** | **1.82** | **[1.04, 2.58]** | **< 0.001** | **0.84** | **[0.79, 0.93]** |
| *Phylloscopus trochilus* | 1.94 | [0.93, 3.05] | 1.63 | [1.13, 2.10] | −0.31 | [−1.27, 0.57] | 0.50 | 0.35 | [0.23, 0.73] |
| *Prinia atrogularis* * | 2.30 | [0.91, 3.72] | 1.55 | [0.95, 2.20] | −0.76 | [−2.06, 0.50] | 0.24 | 0.38 | [0.24, 0.84] |
| *Prunella modularis* | 1.55 | [0.63, 2.47] | 1.69 | [1.36, 2.02] | 0.14 | [−0.75, 0.99] | 0.76 | 0.50 | [0.41, 0.69] |
| *Pyrrhula pyrrhula* * | Excluded (only 1 cuckoo egg sampled) |  |  |  |  |  |  |  |  |
| ***Saxicola rubetra*\*** | **3.97** | **[2.59, 5.36]** | **2.28** | **[1.86, 2.69]** | **−1.68** | **[−3.01, −0.37]** | **0.01** | **0.07** | **[0.03, 0.52]** |
| *Sylvia atricapilla* | 0.93 | [−0.41, 2.23] | 1.31 | [0.60, 2.20] | 0.37 | [−0.57, 1.38] | 0.43 | 0.82 | [0.76, 0.98] |
| *Sylvia borin* | 0.70 | [−0.41, 1.64] | 1.21 | [0.77, 1.62] | 0.51 | [−0.43, 1.40] | 0.28 | 0.74 | [0.67, 0.88] |
| *Sylvia communis* | 0.98 | [−0.58, 2.63] | 1.39 | [0.46, 2.50] | 0.41 | [−0.67, 1.55] | 0.47 | 0.89 | [0.85, 0.99] |
| *Sylvia curruca* * | 1.00 | [−0.62, 2.71] | 1.31 | [0.43, 2.34] | 0.31 | [−0.87, 1.48] | 0.59 | 0.89 | [0.85, 0.99] |
| ***Sylvia hortensis*\*** | **0.25** | **[−1.92, 2.32]** | **1.81** | **[0.58, 3.03]** | **1.57** | **[0.44, 2.79]** | **0.01** | **0.96** | **[0.95, 0.99]** |
| *Sylvia nisoria* * | 1.93 | [0.48, 3.39] | 1.56 | [0.87, 2.38] | −0.36 | [−1.57, 0.81] | 0.56 | 0.52 | [0.36, 0.92] |
| *Troglodytes troglodytes* | 1.10 | [−0.04, 2.28] | 1.36 | [0.86, 1.87] | 0.26 | [−0.73, 1.15] | 0.57 | 0.59 | [0.48, 0.78] |
| *Turdus merula* * | Excluded (only 1 cuckoo egg sampled) |  |  |  |  |  |  |  |  |
| *Turdus philomelos* * |  |  |  |  |  |  |  |  |  |

<sup>†</sup> Here we define cuckoo host-race by their specific host species (i.e., where their eggs were found).

**Table S4.** Data for specimens from which we collected eggshell reflectance. Here we illustrate, for each host species, the number of specimens (eggs), and their respective catalogue numbers at three natural history collections*.

| Latin binomial | No. specimens | Catalogue numbers* |
| --- | --- | --- |
| *Acrocephalus arundinaceus* | 52 | wfvz_151081, wfvz_151077, wfvz_25197, wfvz_151082, wfvz_151078, wfvz_151080, wfvz_25198, wfvz_151079, wfvz_25209, AA10, AA11, AA12, AA13, AA14, AA15, AA16, AA17, AA18, AA19, AA1, AA20, AA21, AA22, AA23, AA24, AA25, AA2, AA3, AA4, AA5, AA6, AA7, AA8, AA9 |
| *Acrocephalus palustris* | 19 | wfvz_23386, wfvz_23388, wfvz_23389, wfvz_23384, wfvz_23390, wfvz_23387 |
| *Acrocephalus schoenobaenus* | 45 | dmnh_27051, dmnh_27052, dmnh_27059, dmnh_27127, dmnh_27128, wfvz_23347, wfvz_23349, wfvz_23348, wfvz_23351, ASC10, ASC11, ASC12, ASC13, ASC14, ASC15, ASC1, ASC2, ASC3, ASC4, ASC5, ASC6, ASC7, ASC8, ASC9 |
| *Acrocephalus scirpaceus* | 52 | wfvz_24897, wfvz_151055, wfvz_24896, wfvz_24909, wfvz_25180, wfvz_25190, wfvz_25188, wfvz_24899, wfvz_24889, wfvz_24915, wfvz_24901, AS10, AS11, AS12, AS13, AS14, AS15, AS16, AS17, AS18, AS19, AS1, AS20, AS21, AS22, AS23, AS24, AS25, AS2, AS3, AS4, AS5, AS6, AS7, AS8, AS9 |
| *Alauda arvensis* | 8 | wfvz_23465, wfvz_151040, wfvz_151034 |
| *Anthus pratensis* | 38 | wfvz_151021, wfvz_151023, wfvz_23343, wfvz_23342, wfvz_23346, wfvz_25254, wfvz_43949, AP11, AP12, AP13, AP14, AP15, AP16, AP17, AP18, AP19, AP20, AP21, AP22, AP23, AP24, AP25, AP26 |
| *Anthus trivialis* | 36 | dmnh_24727, dmnh_26875,<br>dmnh_24919, dmnh_24920,<br>dmnh_24921, dmnh_24924,<br>wfvz_23444, wfvz_113150 |
| *Carduelis cannabina* | 15 | wfvz_23460, wfvz_23458,<br>wfvz_151019, wfvz_43948 |
| *Carduelis chloris* | 1 | dmnh_29452 |
| *Emberiza citrinella* | 16 | wfvz_25669, wfvz_23457,<br>wfvz_23161, wfvz_25668,<br>wfvz_151049 |
| *Emberiza schoeniclus* | 10 | wfvz_151047, wfvz_23394,<br>wfvz_151029 |
| *Erithacus rubecula* | 133 | dmnh_26026, dmnh_26027,<br>dmnh_26028, dmnh_26029,<br>dmnh_26030, dmnh_26031,<br>dmnh_26032, dmnh_26033,<br>dmnh_26034, dmnh_26035,<br>dmnh_26036, dmnh_26037,<br>dmnh_26038, dmnh_26039,<br>dmnh_26040, dmnh_26041,<br>dmnh_26042, dmnh_26043,<br>dmnh_26044, dmnh_26045,<br>dmnh_26046, dmnh_26047,<br>dmnh_26048, dmnh_26049,<br>dmnh_26050, dmnh_26051,<br>dmnh_26052, dmnh_26053,<br>dmnh_26054, dmnh_26055, ER10,<br>ER11, ER12, ER13, ER14, ER15,<br>ER16, ER17, ER18, ER19, ER1,<br>ER20, ER21, ER22, ER23, ER24,<br>ER25, ER26, ER2, ER3, ER4, ER5,<br>ER6, ER7, ER8, ER9 |
| *Fringilla coelebs* | 6 | dmnh_29147, dmnh_29148 |
| *Fringilla montifringilla* | 14 | FM10, FM11, FM12, FM13, FM14,<br>FM1, FM2, FM3, FM4, FM5, FM6,<br>FM7, FM8, FM9 |
| *Lanius collurio* | 52 | dmnh_25534, dmnh_25535,<br>wfvz_25220, wfvz_144265,<br>wfvz_151058, wfvz_25219,<br>wfvz_25217, LC10, LC11, LC12,<br>LC13, LC14, LC15, LC16, LC17,<br>LC18, LC19, LC1, LC20, LC21,<br>LC22, LC23, LC24, LC25, LC2,<br>LC3, LC4, LC5, LC6, LC7, LC8,<br>LC9 |
| *Lanius schach* | 23 | wfvz_43950, wfvz_43954,<br>wfvz_25840, wfvz_110439,<br>wfvz_43953, wfvz_43952,<br>wfvz_43955 |
| *Locustella naevia* | 14 | dmnh_27034, dmnh_27035,<br>dmnh_27045 |
| *Luscinia megarhynchos* | 5 | dmnh_26068 |
| *Motacilla alba* | 80 | dmnh_22784, dmnh_24552,<br>dmnh_24553, dmnh_24555,<br>dmnh_24556, dmnh_24557,<br>dmnh_24558, dmnh_24559,<br>dmnh_24561, wfvz_23227,<br>wfvz_23230, wfvz_23241,<br>wfvz_23239, wfvz_116842, MA10,<br>MA11, MA12, MA13, MA14,<br>MA15, MA16, MA17, MA18,<br>MA19, MA1, MA20, MA21,<br>MA22, MA23, MA24, MA25,<br>MA2, MA3, MA4, MA5, MA6,<br>MA7, MA8, MA9 |
| *Motacilla cinerea* | 8 | dmnh_24518, wfvz_23226 |
| *Motacilla flava* | 13 | dmnh_24482, wfvz_23244,<br>wfvz_23242 |
| *Muscicapa striata* | 13 | wfvz_25250, wfvz_23393,<br>wfvz_25249, wfvz_25251 |
| *Oenanthe oenanthe* | 6 | dmnh_26268 |
| *Passer montanus* | 3 | dmnh_30559 |
| *Phoenicurus phoenicurus* | 44 | dmnh_26165, dmnh_26168,<br>dmnh_26176, wfvz_151039,<br>wfvz_23397, PP10, PP11, PP13,<br>PP14, PP15, PP16, PP17, PP18,<br>PP19, PP1, PP20, PP21, PP22,<br>PP23, PP24, PP25, PP2, PP3, PP4,<br>PP5, PP6, PP7, PP8, PP9 |
| *Phylloscopus trochilus* | 20 | dmnh_27657, dmnh_27658,<br>wfvz_23395, wfvz_151028 |
| *Prinia atrogularis* | 7 | wfvz_13500, wfvz_153526 |
| *Prunella modularis* | 116 | dmnh_25926, dmnh_25927, dmnh_25928, dmnh_25929, dmnh_25930, dmnh_25931, dmnh_25932, dmnh_25933, dmnh_25934, dmnh_25935, dmnh_25936, dmnh_25937, dmnh_25939, dmnh_25940, dmnh_25941, dmnh_25942, dmnh_25943, dmnh_25944, dmnh_25945, dmnh_25946, dmnh_25947, dmnh_25948, dmnh_25950, dmnh_25951, dmnh_25952, dmnh_25953, dmnh_25954, dmnh_25961, dmnh_25962, dmnh_25963, dmnh_25967, dmnh_25969, dmnh_25970, dmnh_25949, PM10, PM11, PM12, PM13, PM14, PM15, PM16, PM17, PM18, PM19, PM1, PM20, PM21, PM22, PM23, PM24, PM25, PM26, PM27, PM2, PM3, PM4, PM5, PM6, PM7, PM8, PM9 |
| *Pyrrhula pyrrhula* | 4 | dmnh_30002 |
| *Saxicola rubetra* | 10 | dmnh_26216, wfvz_23443 |
| *Sylvia atricapilla* | 16 | dmnh_27339, wfvz_25667, wfvz_25665, wfvz_151074, wfvz_151031, wfvz_25666 |
| *Sylvia borin* | 35 | wfvz_23467, wfvz_25670, wfvz_25671, wfvz_23468, SB10, SB11, SB12, SB13, SB14, SB15, SB16, SB17, SB18, SB19, SB1, SB20, SB21, SB22, SB23, SB24, SB25, SB26, SB2, SB3, SB4, SB5, SB6, SB7, SB8, SB9 |
| *Sylvia communis* | 27 | dmnh_27431, dmnh_27432, wfvz_23454, wfvz_151088, wfvz_23453, wfvz_23452, wfvz_187648 |
| *Sylvia curruca* | 10 | dmnh_27472, dmnh_27473, wfvz_23451 |
| *Sylvia hortensis* | 5 | wfvz_151057, wfvz_113245 |
| *Sylvia nisoria* | 7 | wfvz_23456, wfvz_23455 |
| *Troglodytes troglodytes* | 14 | wfvz_151059, wfvz_151060, wfvz_151086, wfvz_25237, wfvz_25231 |

| **Latin binomial** | **No. specimens** | **Catalogue numbers\*** |
| --- | --- | --- |
| *Turdus merula* | 1 | dmnh_26454 |
| *Turdus philomelos* | 2 | dmnh_26726 |

\* catalogue numbers for the Western Foundation of Vertebrate Zoology (wfvz) and the Delaware Museum of Nature and Science (dmnh) include a collection abbreviation before the catalogue number. Egg colour data from Stoddard and Stevens was collected at the Natural History Museum (NHM) and are abbreviated with two letters representing the host species.

